# HRIBO - High-throughput analysis of bacterial ribosome profiling data

**DOI:** 10.1101/2020.04.27.046219

**Authors:** Rick Gelhausen, Florian Heyl, Sarah L. Svensson, Kathrin Froschauer, Lydia Hadjeras, Cynthia M. Sharma, Florian Eggenhofer, Rolf Backofen

**Affiliations:** Bioinformatics Group, Department of Computer Science, University of Freiburg, Georges-Koehler-Allee 106, 79110 Freiburg, Germany; Chair of Molecular Infection Biology II, Institute of Molecular Infection Biology (IMIB), Josef-Schneider-Str. 2 / Bau D15, University of Würzburg, Germany; ZBSA Centre for Biological Systems Analysis, University of Freiburg, Hauptstr. 1, 79104 Freiburg, Germany; Signalling Research Centres BIOSS and CIBSS, University of Freiburg, Schaenzlestr. 18, 79104 Freiburg, Germany

## Abstract

**Motivation:** Ribosome profiling (Ribo-seq) is a powerful approach based on ribosome-protected RNA fragments to explore the translatome of a cell, and is especially useful for the detection of small proteins (<=70 amino acids) that are recalcitrant to biochemical and *in silico* approaches. While pipelines are available to analyze Ribo-seq data, none are designed explicitly for the analysis of Ribo-seq data from prokaryotes, nor are they focused on the discovery of unannotated open reading frames (ORFs) in bacteria.

**Results:** We present `HRIBO` (High-throughput annotation by Ribo-seq), a workflow to enable reproducible and high-throughput analysis of bacterial Ribo-seq data. The workflow performs all required pre-processing and quality control steps. Importantly, `HRIBO` outputs annotation-independent ORF predictions based on two complementary bacteria-focused tools, and integrates them with additional features. This facilitates the rapid discovery of novel ORFs and their prioritization for functional characterization.

**Availability:** `HRIBO` is a free and open source project available under the GPL-3 license at: https://github.com/RickGelhausen/HRIBO

## Introduction

Ribosome profiling (Ribo-seq) is an RNA-seq-based approach that identifies the ribosome-bound fraction of the transcriptome as a proxy for protein expression (Fig 1). Because RNase digestion creates so-called ribosome footprints, it also allows identification of open-reading frame (ORF) boundaries. This makes it remarkably suited to detect small proteins, which are currently underrepresented in genome annotations due to their unique features that preclude conventional biochemical or *in silico* detection [14]. A parallel whole-transcriptome library allows calculation of translation efficiency (TE, the ratio of footprint coverage to transcriptome coverage) and identification of differentially-expressed ORFs. There are existing workflows for the analysis of eukaryotic Ribo-seq data [2], but a specialized solution for bacteria is still missing. Here, we present `HRIBO` (High-throughput annotation by Ribo-seq), a computational pipeline that processes and analyses data from any bacterial Ribo-seq experiment, but also detects translated novel ORFs. This tool is compatible with bacterial annotation and circular genomes, uses an optimized mapping approach suitable for small bacterial genomes, integrates machine learning-based ORF prediction tools designed for/trained on bacterial ORF features and also include two differential expression tools designed for Ribo-seq data. We implemented HRIBO based on snakemake [7], which allows highly reproducible and fully automatic data analysis.

**Figure 1.**
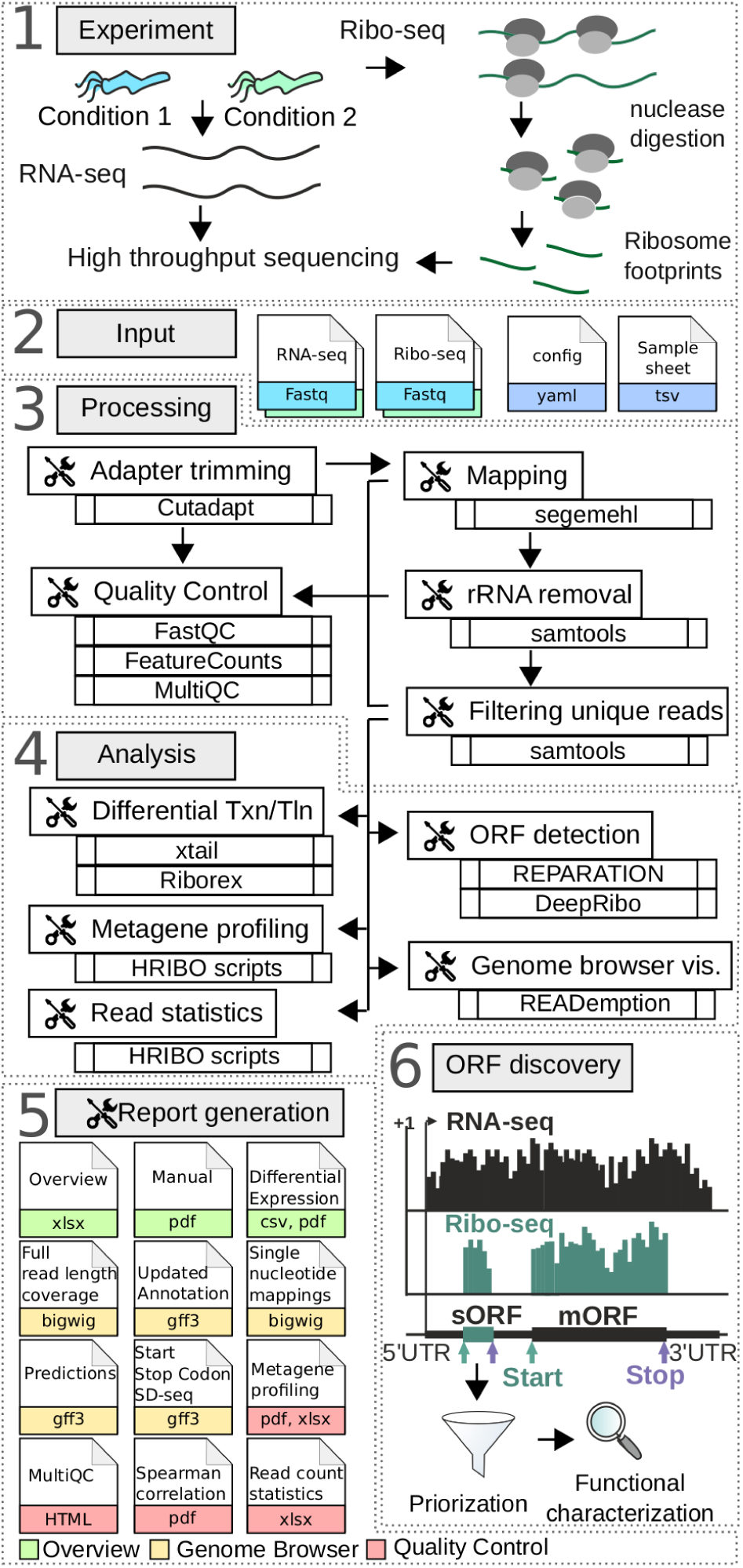
Outline of a bacterial Ribo-seq experiment and data analysis by the `HRIBO` pipeline, (input, processing analysis, output). For details see:https://hribo.readthedocs.io/en/latest.

## Approach

`HRIBO` automatically retrieves tools from bioconda [6] and performs all necessary steps of processing, with pinned and tested tool versions. Detailed documentation, with examples, is available at: https://hribo.readthedocs.io/en/latest/ `HRIBO` (Fig 1, Input) requires the sequencing data files (paired RNA-seq and Ribo-seq libraries for each sample), the genomic sequence/annotation of the organism, and a sample sheet that captures the experimental setup by associating samples/replicates with their experimental condition. As a first step (Fig 1, Processing), `HRIBO` performs adapter trimming with `Cutadapt` [10] and then maps the reads to the genome with `segemehl`, which has higher sensitivity than other mappers [12], but its high computational costs are still acceptable for small genomes.

Multimapping/rRNA reads are then removed before further processing. `FastQC` [1] and `featurecount` [9] reports are created after each processing step and aggregated with `MultiQC` [4], enabling the investigator to identify problems with either the experimental setup (e.g, insufficient rRNA depletion), or the pre-processing (e.g. untrimmed adapters). The resulting mapped reads are then used in subsequent steps (Fig 1, Analysis) to calculate read statistics, detect expressed ORFs (annotated or novel), compute differential expression and coverage tracks, and to perform metagene profiling of global ribosome density at start codons. We have included two complementary tools, both specifically developed for bacterial organisms, to detect ORFs. `REPARATION` [11] uses a random forest-based machine-learning approach. `DeepRibo` [3] is based on convolutional and recurrent neural network approaches. The results of the two prediction tools are aggregated by newly-developed scripts and compiled into a list of detected ORFs. Novel and annotated ORFs that are differentially expressed (transcription and translation) between samples/conditions are detected using the complementary Ribo-seq-specific tools `xtail` [16] and `Riborex` [8]. The ORF list is then enriched with additional computed context information for each ORF, such as length, sequence, normalized read counts, TE, available annotation/novelty, differential expression and which conditions/replicate the ORF was detected in (see Table 1).

**Table 1.**
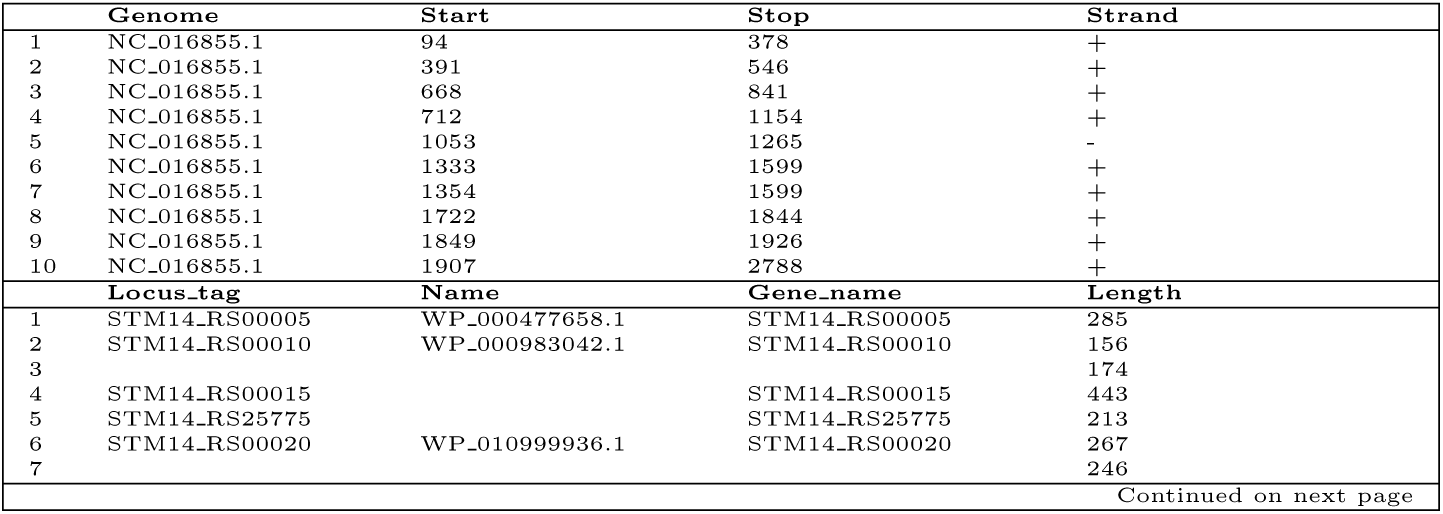
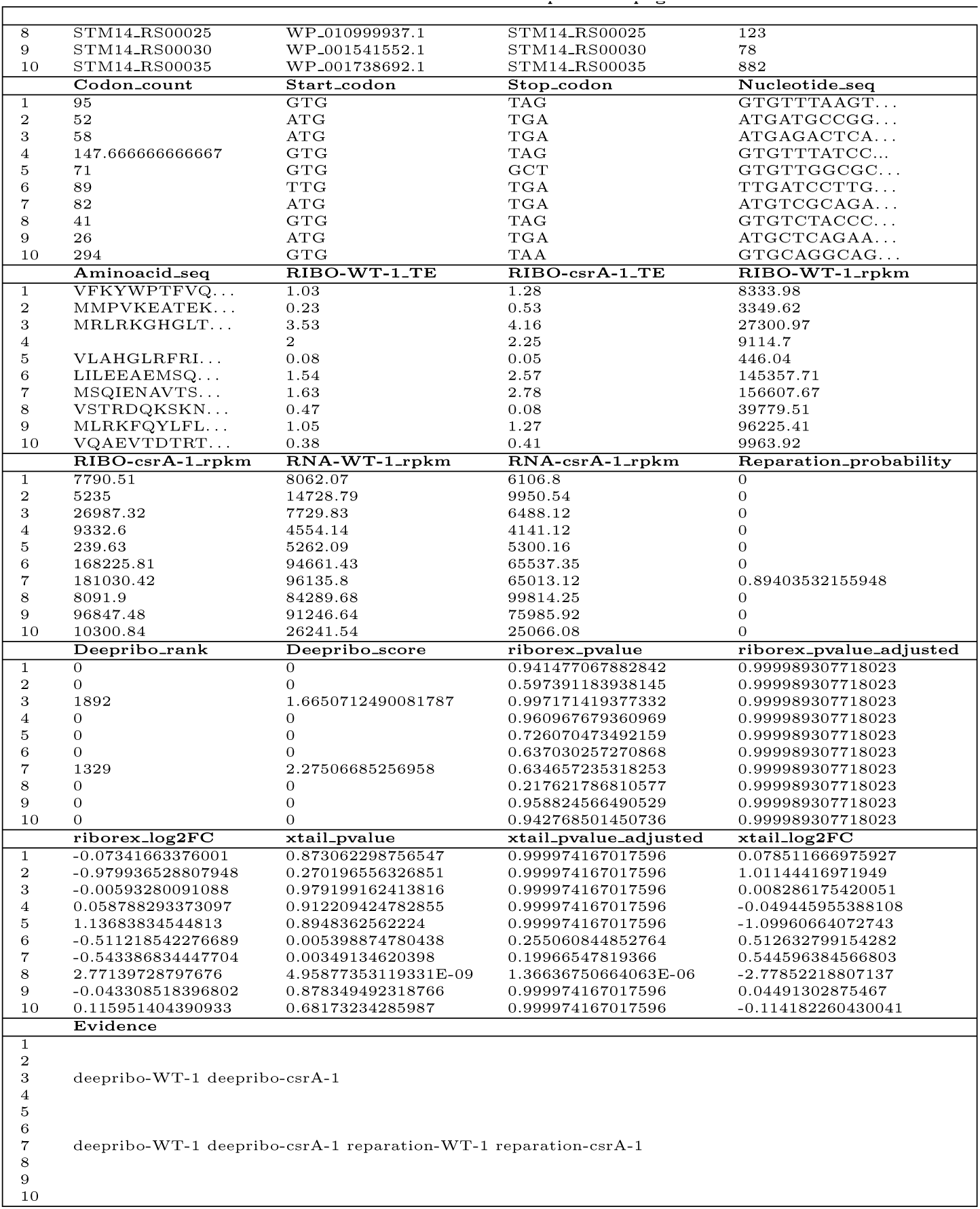
Result overview table, showing the first ten open reading frame entries for the example *Salmonella typhimurium* data set [15] analysed with `HRIBO`. The table contains both annotated and predicted ORFs, from `DeepRibo` and `Reparation` and with context information like length, nucleotide sequence, amino acid sequence, normalized read counts. Moreover the differential transcription and translation is analyzed, with `Riborex` and `xtail`, for all pairwise combinations of provided experimental conditions and added to the table. The evidence column highlights in which experimental condition the ORF was detected by `DeepRibo` and/or `Reparation`. The table, by integrating a plethora of result information enables a stringent screen for candidates for further experimental analysis. For the full output please refer to the complete analyzed dataset referenced in the text.

Interesting candidates can then be inspected by either full-coverage [5] or single nucleotide mapping (5’/3’ end or centre of read) genome browser tracks. Moreover, we developed metagene profiling scripts that analyze a range of read lengths for read density around the start codon (see Figure 2). Finally, the results are collected into a report (Fig 1, Report) that can be easily distributed and contains a detailed manual. An example report for a published dataset [13] can be found here: ftp://biftp.informatik.uni-freiburg.de/pub/HRIBO/HRIBO1.4.0_17-04-20.zip

**Figure 2.**
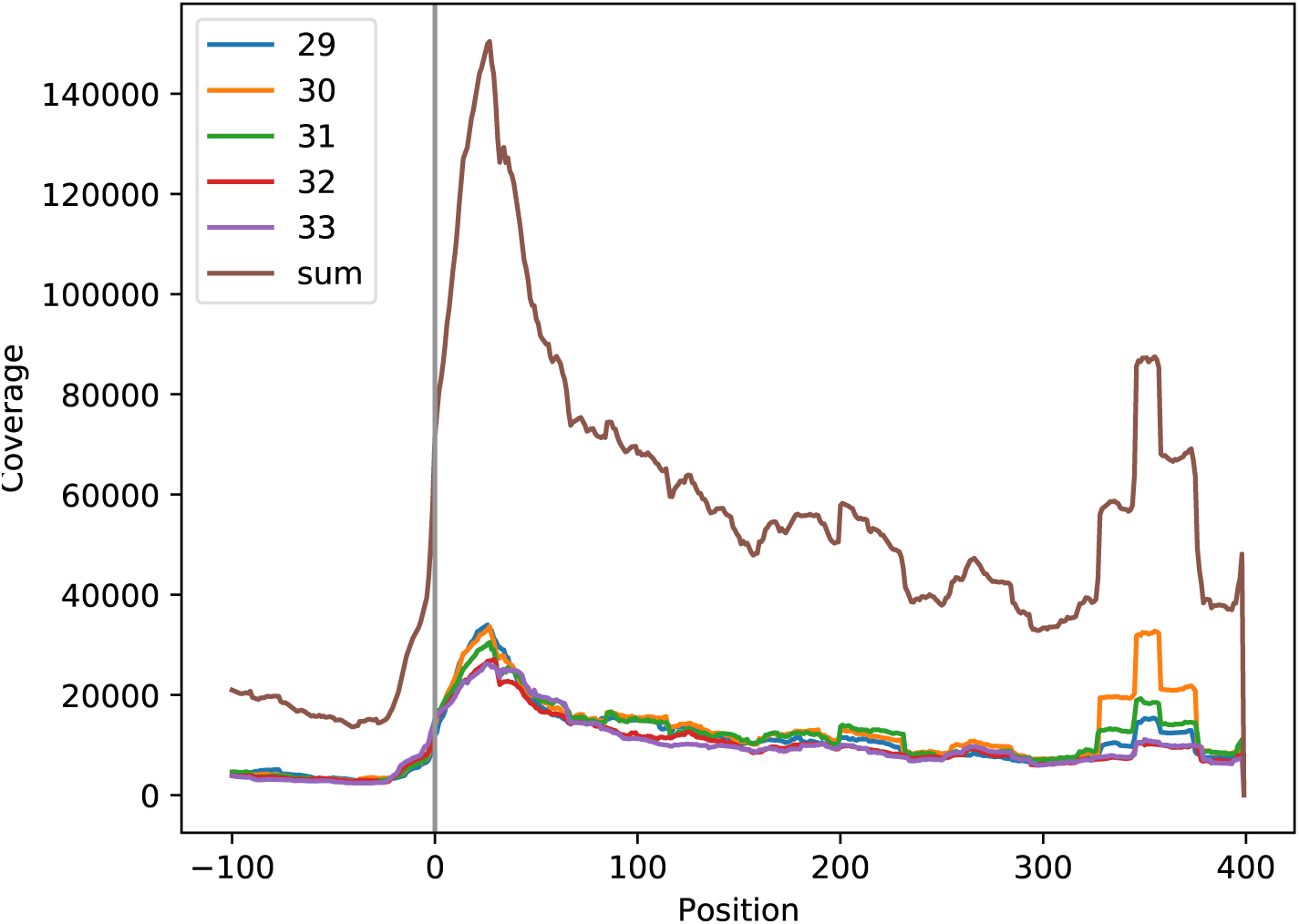
Fragment length specific meta gene profiling from analyzing the example *Salmonella typhimurium* data set with HRIBO version 1.4.0. Metagene profiling is a quality check for Ribo-seq experiments showing the average read distribution around the start codon.

## Conclusion

HRIBO is a reproducible and standardized pipeline to process Ribo-seq data-sets from bacterial organisms, from pre-processing and quality control, to ORF prediction and differential expression analyses. Read information and additional computed features for both annotated and predicted ORFs are summarized, allowing their prioritization for characterization and genome browser tracks for manual inspection. This streamlines the selection of high-confidence, functionally important, translated novel ORFs for further experimental investigation. We plan to add an analysis of translation initiation site data [15] in the future.

## Supporting information

Read the docs - Tool usage guide v1.4.0

Result table from analyzing the example Salmonella typhimurium data set with HRIBO version 1.4.0. The complete report is linked in the manuscript.

Quality report from analyzing the example Salmonella typhimurium data set with HRIBO version 1.4.0. The complete sample is linked in the manuscript.

## Funding

This work was supported by the German Research Foundation (DFG) SCHM 2663/3; the High Performance and Cloud Computing Group, University Tübingen via bwHPC; DFG INST 37/935-1 FUGG. R.G.; DFG grant BA 2168/21-1 SPP 2002; DFG grant 322977937/GRK2344 2017 MeInBio – BioInMe Research Training Group; BMBF “Verbundprojekt RNAProNet, Integration des Netzwerks aus RNA- und Proteinbasierter Regulation - Teilprojekt B - 031L0164B”

---

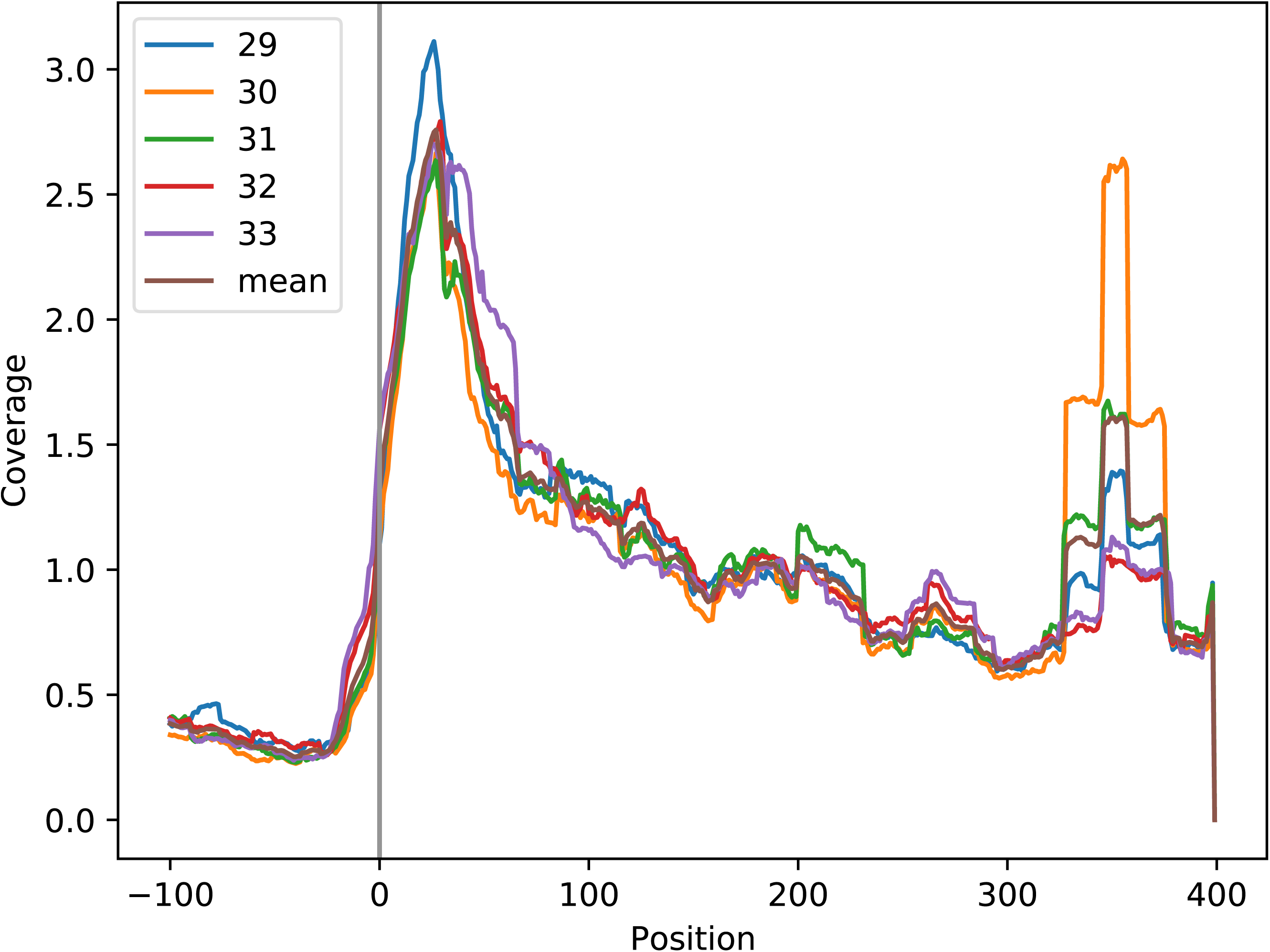

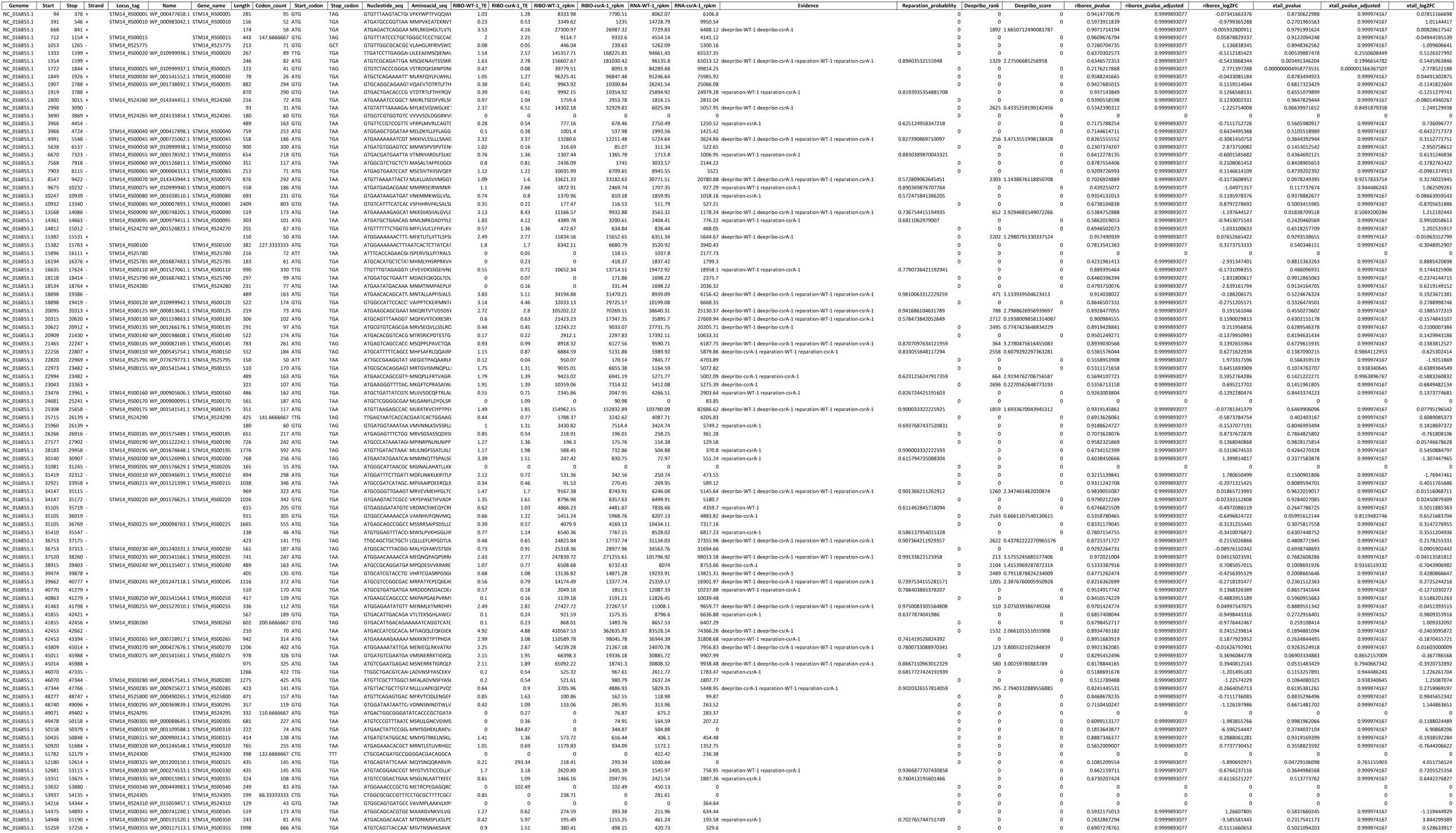

## Notes

### Competing Interest Statement

The authors have declared no competing interest.

