## Supplementary material for "HRIBO - High-throughput analysis of bacterial ribosome profiling data": Read the docs - Tool usage guide v1.4.0

### HRIBO Documentation

***Release 1.4.0***

**Rick Gelhausen**

**Apr 17, 2020**

### DOCUMENTATION

**1 HRIBO** **1**

**4 Example workflow** **23**
4.1 Setup . . . . . . . . . . 23
4.2 Retrieve and prepare input files . . . . . . . . . . 23
4.3 Running the workflow . . . . . . . . . . 26
4.4 Results . . . . . . . . . . 28
4.5 References . . . . . . . . . . 28

**5 Extended workflow** **29**
5.1 Setup . . . . . . . . . . 29
5.2 Retrieve and prepare input files . . . . . . . . . . 29
5.3 Running the workflow . . . . . . . . . . 32
5.4 Results . . . . . . . . . . 34
5.5 References . . . . . . . . . . 35

**6 Frequently asked questions** **37**

6.1 Q: When using singularity I get `ERROR : Failed to set effective UID to 0` . . . . . . 37

**Bibliography** **39**

CHAPTER

ONE

### HRIBO

#### 1.1 Introduction

HRIBO is a workflow for the analysis of prokaryotic Ribo-Seq data. HRIBO is available on github. It includes among others, prediction of novel open reading frames (**ORFs**), metagene profiling, quality control and differential expression analysis. The workflow is based on the workflow management system **snakemake** and handles installation of all dependencies via bioconda [GruningDSjodin+17] and docker, as well as all processings steps. The source code of HRIBO is open source and available under the License **GNU General Public License 3**. Installation and basic usage is described below.

**Note:** For a detailed step by step tutorial on how to use this workflow on a sample dataset, please refer to our *example-workflow*.

#### 1.2 Program flowchart

The following flowchart describes the processing steps of the workflow and how they are connected:

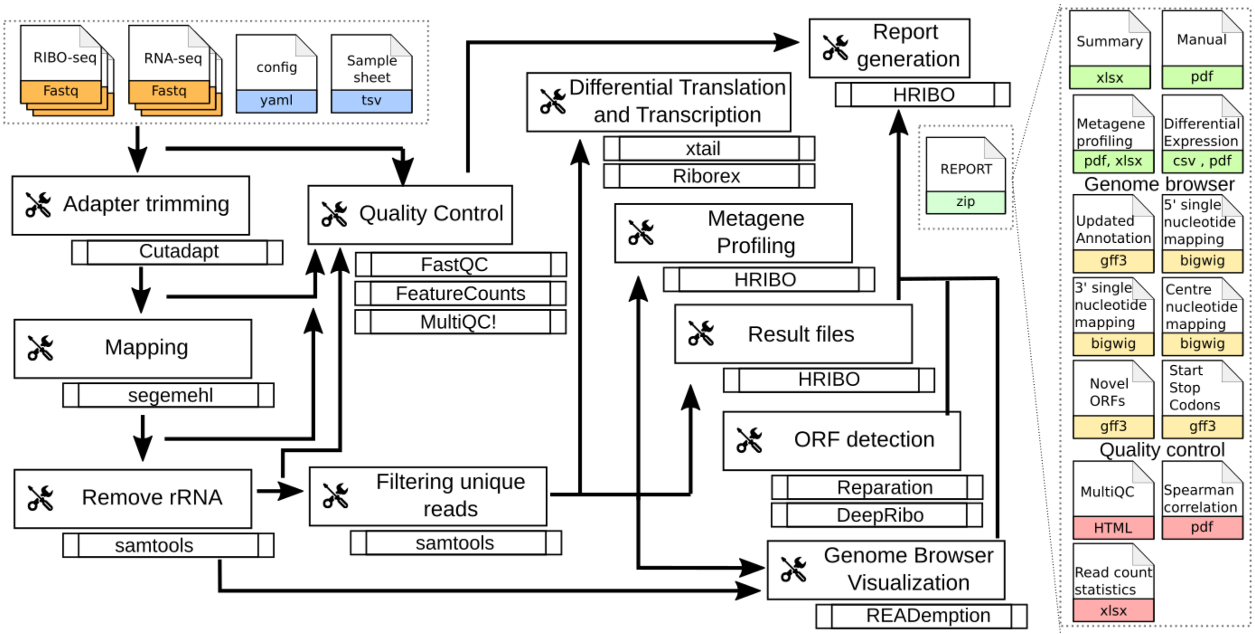

#### 1.3 Requirements

In the following, we describe all the required files and tools needed to run our workflow.

#### 1.4 Tools

##### 1.4.1 miniconda3

As this workflow is based on the workflow management system snakemake [KosterR18], **Snakemake** will download all necessary dependencies via conda.

We strongly recommend installing miniconda3 with `python3.7`.

After downloading the `miniconda3` version suiting your linux system, execute the downloaded bash file and follow the instructions given.

##### 1.4.2 snakemake

**Note:** HRIBO requires `snakemake (version>=5.5.1)`

The newest version of `snakemake` as well as the `squashfs-tools` required for `singularity` can be downloaded via conda using the following command:

```
conda create -c conda-forge -c bioconda -n snakemake snakemake squashfs-tools
```

This creates a new conda environment called `snakemake` and installs `snakemake` into the environment. The environment can be activated using:

```
conda activate snakemake
```

and deactivated using:

```
conda deactivate
```

##### 1.4.3 singularity

**Warning:** This dependency is only required if you intend to use the prediction tool `deepribo`. The rest of the workflow does not require `singularity`. `deepribo` is deactivated by default. For more details on activating `deepribo`, please refer to *Activating DeepRibo*.

In order to support docker container, snakemake requires singularity. This is used to retrieve tools that are not available on conda as of now.

An in-depth installation tutorial for singularity can be found on the singularity webpage.

**Note:** we strongly suggest to install the newest version of singularity. We tested our worklow on `singularity v3.4.2`.

##### 1.4.4 HRIBO

Using the workflow requires `HRIBO`. The latest version is available on our GitHub page.

In order to run the workflow, we suggest that you download the `HRIBO` into your project directory. The following command creates an example directory and changes into it:

```
mkdir project
cd project
```

Now, download and unpack the latest version of `HRIBO` by entering the following commands:

```
wget https://github.com/RickGelhausen/HRIBO/archive/1.4.0.tar.gz
tar -xzf 1.4.0.tar.gz; mv HRIBO-1.4.0 HRIBO; rm 1.4.0.tar.gz;
```

`HRIBO` is now in a subdirectory of your project directory.

#### 1.5 Input files

Several input files are required in order to run the workflow, a genome file (`.fa`), an annotation file (`.gff/.gtf`) and compressed sequencing files (`.fastq.gz`).

| File name | Description |
|---|---|
| annotation.gff | user-provided annotation file with genomic features |
| genome.fa | user-provided genome file containing the genome sequence |
| <method>-<conditon>-<replicate>.fastq.gz | user-provided compressed sequencing files |
| config.yaml | configuration file to customize the workflow |
| samples.tsv | sample file describing the relation between the input fastq files |

##### 1.5.1 annotation.gff and genome.fa

We recommend retrieving both the genome and the annotation files for your organism from National Center for Biotechnology Information (NCBI) or Ensembl Genomes [ZAA+18].

> **Warning:** if you use custom annotation files, ensure that you adhere to the gtf/gff standard. Wrongly formatted files are likely to cause problems with downstream tools.

**Note:** For detailed information about downloading and unpacking these files, please refer to our *example-workflow*.

##### 1.5.2 input .fastq files

These are the input files provided by the user. Both single end and paired end data is supported.

**Note:** As most downstream tools do not support paired end data, we combine the paired end data into single end data using flash2 . For more information about how to use paired-end data please refer to the *workflow-configuration*.

**Note:** Please ensure that you compress your files in `.gz` format.

Please ensure that you move all input `.fastq.gz` files into a folder called **fastq** (Located in your project folder):

```
mkdir fastq
cp *.fastq.gz fastq/
```

##### 1.5.3 Sample sheet and configuration file

In order to run `HRIBO`, you have to provide a sample sheet and a configuration file. There are templates for both files available in the `HRIBO` folder, in the subfolder `templates`. The configuration file is used to allow the user to easily customize certain settings, like the adapter sequence. The sample sheet is used to specify the relation of the input `.fastq` files (condition / replicate etc. . . )

Copy the templates of the sample sheet and the configuration file into the `HRIBO` folder:

```
cp HRIBO/templates/samples.tsv HRIBO/
cp HRIBO/templates/config.yaml HRIBO/
```

Customize the `config.yaml` using your preferred editor. It contains the following variables:

- **adapter:** specify the adapter sequence to be used.
- **samples:** the location of the samples sheet created in the previous step.
- **alternativestartcodons:** specify a comma separated list of alternative start codons.
- **differentialexpression:** specify whether you want to activate differential expresssion analysis. ("yes/no")
- **deepribo:** specify whether you want to activate deepribo ORF prediction. ("yes/no")

Edit the sample sheet corresponding to your project. It contains the following variables:

- **method:** indicates the method used for this project, here RIBO for ribosome profiling and RNA for RNA-seq.
- **condition:** indicates the applied condition (e.g. A, B, . . . ).
- **replicate:** ID used to distinguish between the different replicates (e.g. 1,2, . . . )
- **inputFile:** indicates the according fastq file for a given sample.

**Note:** If you have paired end data, please ensure that you use the `samples_pairedend.tsv` file.

As seen in the `samples.tsv` template:

| method | condition | replicate | fastqFile |
|---|---|---|---|
| RIBO | A | 1 | fastq/RIBO-A-1.fastq.gz |
| RIBO | A | 2 | fastq/RIBO-A-2.fastq.gz |
| RIBO | B | 1 | fastq/RIBO-B-1.fastq.gz |
| RIBO | B | 2 | fastq/RIBO-B-2.fastq.gz |
| RNA | A | 1 | fastq/RNA-A-1.fastq.gz |
| RNA | A | 2 | fastq/RNA-A-2.fastq.gz |
| RNA | B | 1 | fastq/RNA-B-1.fastq.gz |
| RNA | B | 2 | fastq/RNA-B-2.fastq.gz |

**Note:** This is just an example, please refer to our *example-workflow* for another example.

##### 1.5.4 cluster.yaml

In the `HRIBO` folder, we provide two `<cluster>.yaml` files needed by snakemake in order to run on a cluster system:

- **sge.yaml** - for grid based queuing systems
- **torque.yaml** - for torque based queuing systems

#### 1.6 Output files

In the following tables all important output files of the workflow are listed.

**Note:** Files create as intermediate steps of the workflow are omitted from this list. (e.g. `.bam` files)

**Note:** For more details about the output files, please refer to the *analysis results*.

##### 1.6.1 Single-file Output

| File name | Description |
|---|---|
| samples.xlsx | Excel version of the input samples file. |
| manual.pdf | A PDF file describing the analysis. |
| annotation_total.xlsx | Excel file containing detailed measures for every feature in the input annotation using read counts containing multi-mapping reads. |
| annotation_unique.xlsx | Excel file containing detailed measures for every feature in the input annotation using read counts containing no multi-mapping reads. |
| total_read_counts.xlsx | Excel file containing read counts with multi-mapping reads. |
| unique_read_counts.xlsx | Excel file containing read counts without multi-mapping reads. |
| multiqc_report.html | Quality control report combining all finding of individual fastQC reports into a well structured overview file. |
| heatmap_SpearmanCorr_readCounts.pdf | PDF file showing the Spearman correlation between all samples. |
| predictions_reparation.xlsx | Excel file containing detailed measures for every ORF detected by reparation. |
| predictions_reparation.gff | GFF file containing ORFs detected by reparation, for genome browser visualization. |
| potentialStartCodons.gff | GFF file for genome browser visualization containing all potential start codons in the input genome. |
| potentialStopCodons.gff | GFF file for genome browser visualization containing all potential stop codons in the input genome. |
| potentialRibosomeBindingSite.gff | GFF file for genome browser visualization containing all potential ribosome binding sites in the input genome. |
| potentialAlternativeStartCodons.gff | GFF file for genome browser visualization containing all potential alternative start codons in the input genome. |

##### 1.6.2 Multi-file Output

| File name | Description |
|---|---|
| riborex/<contrast>_sorted.csv | Differential expression results by Riborex, sorted by pvalue. |
| riborex/<contrast>_significant.csv | Differential expression results by Riborex, only significant results. (pvalue < 0.05) |
| xtail/<contrast>_sorted.csv | Differential expression results by xtail, sorted by pvalue. |
| xtail/<contrast>_significant.csv | Differential expression results by xtail, only significant results. (pvalue < 0.05) |
| xtail/r_<contrast>.pdf | Differential expression results by xtail, plot with RPF-to-mRNA ratios. |
| xtail/fc_<contrast>.pdf | Differential expression results by xtail, plot with log2 fold change of both mRNA and RPF. |
| <method>-<condition>-<replicate>.X.Y.Z.bw | BigWig file for genome browser visualization, containing a single nucleotide mapping around certain regions. |
| <accession>_Z.Y_profiling.xlsx/tsv | Excel and tsv files containing raw data of the metagene analysis. |
| <accession>_Z.Y_profiling.pdf | visualization of the metagene analysis. |

**Note:** <contrast> represents a pair of conditions that are being compared.

**Note:** The BigWig files are available for different normalization methods, strands and regions, X=(min/mil) Y=(forward/reverse) Z=(fiveprime, threeprime, global, centered).

#### 1.7 Tool Parameters

The tools used in our workflow are listed below, with links to their respective webpage and a short description.

| Tool | Version | Special parameters used |
|---|---|---|
| cutadapt | 2.1 | Adapter removal and quality trimming |
| fastQC | 0.11.9 | Quality control |
| multiQC | 1.8 | Quality control report |
| segemehl | 0.3.4 | Mapping of reads |
| flash2 | 2.2.00 | Merging paired end samples into single end |
| cufflinks | 2.2.1 | Used to convert gff to gtf |
| bedtools | 2.27.1 | Collection of useful processing tools (e.g. read counting etc...) |
| reparation_blast | 1.0.9 | Prediction of novel Open Reading frames |
| deepribo | 1.1 | Prediction of novel Open Reading frames |
| riborex | 2.4.0 | Differential expression analysis |
| xtail | 1.1.5 | Differential expression analysis |

#### 1.8 Report

In order to aggregate the final results into a single folder structure and receive a date-tagged `.zip` file, you can use the `makereport.sh` script.

```
bash HRIBO/scripts/makereport.sh <reportname>
```

**Note:** Examples of how this output can look are available here .

#### 1.9 Example-workflow

A detailed step by step tutorial is available at: *example-workflow*.

#### 1.10 References

CHAPTER

### TWO

### WORKFLOW CONFIGURATION

This workflow allows different customization to be able to handle different types of input data. On this page we explain the different options that can be set to easily customize the workflow.

#### 2.1 Default workflow

In order to explain what customizations are possible, we will first have a look at the default workflow.

Default:

- Single-end fastq files
- Differential expression analysis: on
- DeepRibo predictions: off

For the default workflow, we expect the `.fastq` files to be in single-end format. Additionally, we activated differential expression by default. Differential expression requires multiple conditions and RIBO and RNA samples. A possible `sample.tsv` would look as follows:

| method | condition | replicate | fastqFile |
|---|---|---|---|
| RIBO | A | 1 | fastq/RIBO-A-1.fastq.gz |
| RIBO | A | 2 | fastq/RIBO-A-2.fastq.gz |
| RIBO | B | 1 | fastq/RIBO-B-1.fastq.gz |
| RIBO | B | 2 | fastq/RIBO-B-2.fastq.gz |
| RNA | A | 1 | fastq/RNA-A-1.fastq.gz |
| RNA | A | 2 | fastq/RNA-A-2.fastq.gz |
| RNA | B | 1 | fastq/RNA-B-1.fastq.gz |
| RNA | B | 2 | fastq/RNA-B-2.fastq.gz |

**Note:** By default only reparation predictions are used. The reason for this is that DeepRibo is not available on conda as of now and therfore requires additional tweaks to run it. The process is explained *below*.

#### 2.2 No differential expression

If you do not have multiple conditions and differential expression is activated, you will receive an error message. To deactivate differential expression, you have to edit the `config.yaml` file.

```
adapter: ""
samples: "HRIBO/samples.tsv"
alternativestartcodons: "GTG,TTG"
# Differential expression: on / off
differentialexpression: "off"
# Deepribo predictions: on / off
deepribo: "off"
```

This will allow you the use of a sample.tsv like:

| method | condition | replicate | fastqFile |
|---|---|---|---|
| RIBO | A | 1 | fastq/RIBO-A-1.fastq.gz |
| RIBO | A | 2 | fastq/RIBO-A-2.fastq.gz |
| RNA | A | 1 | fastq/RNA-A-1.fastq.gz |
| RNA | A | 2 | fastq/RNA-A-2.fastq.gz |

#### 2.3 Activating DeepRibo

Activating DeepRibo predictions will give you a different file with ORF predictions. By experience, the top DeepRibo results tend to be better than those of reparation. For archea, where reparation performs very poorly, DeepRibo is the preferred option.

**Note:** In order to use DeepRibo, the tool `singularity` is required. Please refer to the *overview* for details on the installation.

Once you have installed `singularity` turn on DeepRibo in the `config.yaml`:

```
adapter: ""
samples: "HRIBO/samples.tsv"
alternativestartcodons: "GTG,TTG"
# Differential expression: on / off
differentialexpression: "on"
# Deepribo predictions: on / off
deepribo: "on"
```

When calling snakemake, you will now require additional commandline arguments:

- **–use-singularity:** specify that snakemake can now download and use docker container via singularity.
- **–singularity-args " -c ":** specify the `--contain` option to ensure that only the docker containers file system will be used.

**Warning:** DeepRibo cannot cope with genomes containing special `IUPAC symbols`, ensure that your genome file contains only `A`, `G`, `C`, `T`, `N` symbols.

##### 2.3.1 If you run deepribo locally

When running the workflow with DeepRibo locally it might be advised to additionally use the `--greediness 0` option, if you do not have a lot of cores available locally. This will cause the workflow to submit fewer jobs at the same time. This especially important for DeepRibo as we observed that a single DeepRibo job can finish in less than an hour if it does not have to fight for cores with another DeepRibo job. Otherwise, it can run for several hours at a time.

```
snakemake --use-conda --use-singularity --singularity-args " -c " -s HRIBO/Snakefile -
↪-configfile HRIBO/config.yaml --directory ${PWD} -j 10 --latency-wait 60
```

##### 2.3.2 If you run deepribo on a cluster system

When running the workflow with DeepRibo on a cluster system. You have to add the above commandline arguments to your submission script.

```
#!/bin/bash
#PBS -N <ProjectName>
#PBS -S /bin/bash
#PBS -q "long"
#PBS -d <PATH/ProjectFolder>
#PBS -l nodes=1:ppn=1
#PBS -o <PATH/ProjectFolder>
#PBS -j oe
cd <PATH/ProjectFolder>
source activate HRIBO
snakemake --latency-wait 600 --use-conda --use-singularity --singularity-args " -c " -
↪s HRIBO/Snakefile --configfile HRIBO/config.yaml --directory ${PWD} -j 20 --cluster-
↪config HRIBO/templates/torque-cluster.yaml --cluster "qsub -N {cluster.jobname} -S /
↪bin/bash -q {cluster.qname} -d <PATH/ProjectFolder> -l {cluster.resources} -o
↪{cluster.logoutputdir} -j oe"
```

**Note:** If you cannot install `singularity` on your cluster, check whether there are modules available for you cluster system.

You can then create an additional submission script that will tell snakemake to activate the module before running jobs. An example of this would look as follows:

`jobscript.sh`

```
#!/bin/bash
module load devel/singularity/3.4.2
# properties = {properties}
{exec_job}
```

Then add the jobscript to the snakemake call:

```
#!/bin/bash
#PBS -N <ProjectName>
#PBS -S /bin/bash
#PBS -q "long"
#PBS -d <PATH/ProjectFolder>
#PBS -l nodes=1:ppn=1
#PBS -o <PATH/ProjectFolder>
```

(continues on next page)

(continued from previous page)

```
#PBS -j oe
cd <PATH/ProjectFolder>
source activate HRIBO
snakemake --latency-wait 600 --use-conda --use-singularity --singularity-args " -c " -
↪-jobscript jobscript.sh -s HRIBO/Snakefile --configfile HRIBO/config.yaml --
↪directory ${PWD} -j 20 --cluster-config HRIBO/templates/torque-cluster.yaml --
↪cluster "qsub -N {cluster.jobname} -S /bin/bash -q {cluster.qname} -d <PATH/
↪ProjectFolder> -l {cluster.resources} -o {cluster.logoutputdir} -j oe"
```

This will specify to snakemake that it will execute `module load devel/singularity/3.4.2` when submitting each job.

**Warning:** This is a specific example for our TORQUE cluster system. The specific way of loading modules, as well as the available modules, can differ on each system.

#### 2.4 Paired-end support

We allow paired-end data in our workflow. Unfortunately, many of the downstream tools, like the prediction tools, cannot use paired-end data. Therefore, we use the tool `flash2` to convert paired-end data to single-end data.

In order to use paired-end data, simply replace the `Snakefile` with the `Snakefile_pairedend`. This will now require a special `samples_pairedend.tsv`, which is also available in the HRIBO templates folder.

| method | condition | replicate | fastqFile1 | fastqFile2 |
|---|---|---|---|---|
| RIBO | A | 1 | fastq/RIBO-A-1_R1.fastq.gz | fastq/RIBO-A-1_R2.fastq.gz |
| RIBO | A | 2 | fastq/RIBO-A-2_R1.fastq.gz | fastq/RIBO-A-2_R2.fastq.gz |
| RIBO | B | 1 | fastq/RIBO-B-1_R1.fastq.gz | fastq/RIBO-B-1_R2.fastq.gz |
| RIBO | B | 2 | fastq/RIBO-B-2_R1.fastq.gz | fastq/RIBO-B-2_R2.fastq.gz |
| RNA | A | 1 | fastq/RNA-A-1_R1.fastq.gz | fastq/RNA-A-1_R2.fastq.gz |
| RNA | A | 2 | fastq/RNA-A-2_R1.fastq.gz | fastq/RNA-A-2_R2.fastq.gz |
| RNA | B | 1 | fastq/RNA-B-1_R1.fastq.gz | fastq/RNA-A-1_R2.fastq.gz |
| RNA | B | 2 | fastq/RNA-B-2_R1.fastq.gz | fastq/RNA-A-1_R2.fastq.gz |

CHAPTER

THREE

### ANALYSIS RESULT FILES

The important files in this workflow are listed and explained below.

#### 3.1 ORF Predictions

The output files containing information about predicted Open Reading Frames, these also contain novel predictions.

##### 3.1.1 predictions_reparation.xlsx

This file contains all `reparation` ORF predictions.

| Column name | Description |
|---|---|
| Genome | The genome accession identifier. |
| Source | The source of the ORF. (here reparation) |
| Feature | The feature of the ORF (here CDS) |
| Start | The start position of the ORF. |
| Stop | The stop position of the ORF. |
| Strand | The strand of the ORF. (+/-) |
| Locus_tag | If the detected ORF is already in the annotation, this gives its locus tag. |
| Name | If the detected ORF is already in the annotation, this gives its name. |
| Length | The length of the ORF. |
| Codon_count | The number of codons in the ORF. (length/3) |
| <method>-<condition>-<replicate>_TE | The translational efficiency for the given sample. |
| <method>-<condition>-<replicate>_rpkm | The RPKM for the given sample. |
| Evidence | The <condition>-<replicate> sample in which this ORF was predicted. |
| Start_codon | The start codon of the ORF. |
| Stop_codon | The stop codon of the ORF. |
| Nucleotide_seq | The nucleotide sequence of the ORF. |
| Aminoacid_seq | The amino acid sequence of the ORF. |
| Product | only available in annotation_X.xlsx (X=total/unique) |
| Note | only available in annotation_X.xlsx (X=total/unique) |

##### 3.1.2 predictions_reparation.gff

An annotation file in `.gff3` format containing all predictions of `reparation` for visualization in a genome browser.

##### 3.1.3 predictions_deepribo.xlsx

**Note:** These files are only available when activating DeepRibo predictions in the `config.yaml`. (see workflow-configuration <workflow-configuration:workflow-configuration>)

This file contains all `DeepRibo` ORF predictions.

| Column name | Description |
|---|---|
| Genome | The genome accession identifier. |
| Source | The source of the ORF. (here reparation) |
| Feature | The feature of the ORF (here CDS) |
| Start | The start position of the ORF. |
| Stop | The stop position of the ORF. |
| Strand | The strand of the ORF. (+/-) |
| Pred_value | The value DeepRibo attributes the given prediction. |
| Pred_rank | The rank calculated from the prediction value. (the best prediction has rank 1) |
| Novel_rank | A special ranking involving only novel ORFs that are not in the annotation. |
| Locus_tag | If the detected ORF is already in the annotation, this gives its locus tag. |
| Name | If the detected ORF is already in the annotation, this gives its name. |
| Length | The length of the ORF. |
| Codon_count | The number of codons in the ORF. (length/3) |
| <method>-<condition>-<replicate>_TE | The translational efficiency for the given sample. |
| <method>-<condition>-<replicate>_rpkm | The RPKM for the given sample. |
| Evidence | The <condition>-<replicate> sample in which this ORF was predicted. |
| Start_codon | The start codon of the ORF. |
| Stop_codon | The stop codon of the ORF. |
| Nucleotide_seq | The nucleotide sequence of the ORF. |
| Aminoacid_seq | The amino acid sequence of the ORF. |
| Product | only available in annotation_X.xlsx (X=total/unique) |
| Note | only available in annotation_X.xlsx (X=total/unique) |

##### 3.1.4 predictions_deepribo.gff

**Note:** These files are only available when activating DeepRibo predictions in the `config.yaml`. (see workflow-configuration <workflow-configuration:workflow-configuration>)

An annotation file in `.gff3` format containing all predictions of *DeepRibo* for visualization in a genome browser.

#### 3.2 Quality control

This comprises all files that can help to perform quality control on all input samples.

##### 3.2.1 multiqc_report.html

The multiQC report collects information from different tools, including `fastQC` and `subread featurecounts`. The general statistics give an overview over:

- the number of duplicates
- the GC content
- the average read lengths
- the number of reads (in millions)

These statistics are collected after each processing step of our pipeline.

- **raw:** the unprocessed data
- **trimmed:** the data after trimming the adapter sequences
- **mapped:** the data after mapping with Segemehl
- **unique:** the data after removing multi-mapping reads
- **norRNA:** the data after filtering out the rRNA

Further, feature counts are provided for different features from the annotation file. (i.e. how many reads map to each feature) This includes, all(featurecount), rRNA, norRNA(after filtering), tRNA and ncRNA. Following is a fastQC report including sequence counts, sequence quality histograms, per sequence quality scores, per base sequence content, per sequence GC content, per base N content, sequence length distribution, sequence duplication levels, overrepresented features, adapter content and a status overview.

##### 3.2.2 heatmap_SpearmanCorr_readCounts.pdf

Spearman correlation coefficients of read counts. The dendrogram indicates which samples read counts are most similar to each other. Since there should be always a higher correlation between experiments with the same condition and experiment type (e.g. replicates) and not others, this is a rapid way to quality-control the labeling/consistency of input data.

##### 3.2.3 annotation_total.xlsx

This file contains detailed measures for every feature in the input annotation using read counts including multi-mapping reads.

| Column name | Description |
| --- | --- |
| Genome | The genome accession identifier. |
| Source | The source of the annotated feature. |
| Feature | The feature of the annotated feature. |
| Start | The start position of the annotated feature. |
| Stop | The stop position of the annotated feature. |
| Strand | The strand of the annotated feature. (+/-) |
| Locus_tag | The locus tag of the annotated feature. (if available) |
| Name | The name of the annotated feature. (if available) |
| Length | The length of the annotated feature. |
| Codon_count | The number of codons in the annotated feature. (length / 3) |
| <method>-<condition>-<replicate>_TE | The translational efficiency for the given sample. |
| <method>-<condition>-<replicate>_rpkm | The RPKM for the given sample. (ReadsPerKilobaseMillion) |
| Evidence | only available for predicted ORFs |
| Start_codon | The start codon of the annotated feature. |
| Stop_codon | The stop codon of the annotated feature. |
| Nucleotide_seq | The nucleotide sequence of the annotated feature. |
| Aminoacid_seq | The amino acid sequence of the annotated feature. |
| Product | The product of the annotated feature. (if available) |
| Note | The note of the annotated feature. (if available) |

##### 3.2.4 total_read_counts.xlsx

This file shows the overall read-counts for each feature annotated in the user-provided annotation, after mapping and before removal of multi-mapping reads.

##### 3.2.5 annotation_unique.xlsx

This file contains detailed measures for every feature in the input annotation using read counts after removal of multi-mapping reads.

| Column name | Description |
|---|---|
| Genome | The genome accession identifier. |
| Source | The source of the annotated feature. |
| Feature | The feature of the annotated feature. |
| Start | The start position of the annotated feature. |
| Stop | The stop position of the annotated feature. |
| Strand | The strand of the annotated feature. (+/-) |
| Locus_tag | The locus tag of the annotated feature. (if available) |
| Name | The name of the annotated feature. (if available) |
| Length | The length of the annotated feature. |
| Codon_count | The number of codons in the annotated feature. (length / 3) |
| <method>-<condition>-<replicate>_TE | The translational efficiency for the given sample. |
| <method>-<condition>-<replicate>_rpkm | The RPKM for the given sample. (ReadsPerKilobaseMillion) |
| Evidence | only available for predicted ORFs |
| Start_codon | The start codon of the annotated feature. |
| Stop_codon | The stop codon of the annotated feature. |
| Nucleotide_seq | The nucleotide sequence of the annotated feature. |
| Aminoacid_seq | The amino acid sequence of the annotated feature. |
| Product | The product of the annotated feature. (if available) |
| Note | The note of the annotated feature. (if available) |

##### 3.2.6 unique_read_counts.xlsx

This file shows the overall read-counts for each feature annotated in the user-provided annotation, after mapping and after removal of multi-mapping reads.

#### 3.3 genome-browser

The files that can be used for visualization in a genome browser.

##### 3.3.1 updated_annotation.gff

A gff track containing both the original annotation together with the new predictions by reparation.

##### 3.3.2 potentialStartCodons.gff

A genome browser track with all possible start codons.

##### 3.3.3 potentialStopCodons.gff

A genome browser track with all possible stop codons.

##### 3.3.4 potentialRibosomeBindingSite.gff

A genome browser track with possible ribosome binding sites.

##### 3.3.5 potentialAlternativeStartCodons.gff

A genome browser track with alternative start codons.

##### 3.3.6 BigWig coverage files

We offer many different single nucleotide mapping bigwig files for genome browser visualization. These files are available for different regions and performed with different methods.

- **global:** full read is mapped
- **centered:** region around the center.
- **threeprime:** region around the three prime end.
- **fiveprime:** region around the five prime end.

These are all available with the following processing methods:

- **raw:** raw, unprocessed files
- **min:** normalized with by number of minimal total reads per sample (factor = min. number of reads / number of reads)
- **mil:** normalized with by 1000000 (factor = 1000000 / number of reads)

#### 3.4 Differential Expression

Files related to the differential expression analysis.

##### 3.4.1 riborex/<contrast>_sorted.xlsx

Table containing all differential expression results from *riborex*.

##### 3.4.2 riborex/<contrast>_significant.xlsx

Table containing significant differential expression results from *riborex* (pvalue < 0.05).

##### 3.4.3 xtail/<contrast>_sorted.xlsx

Table containing all differential expression results from *xtail*.

##### 3.4.4 xtail/<contrast>_significant.xlsx

Table containing significant differential expression results from *xtail* (pvalue < 0.05).

##### 3.4.5 xtail/r_<contrast>.pdf

This figure shows the RPF-to-mRNA ratios in two conditions, where the position of each gene is determined by its RPF-to-mRNA ratio (log2R) in two conditions, represented on the x-axis and y-axis respectively. The points will be color-coded with the pvalue final obtained with xtail (more significant p values having darker color)

- **blue:** for genes with log2R larger in first condition than second condition.
- **red:** for genes with log2R larger in second condition than the first condition.
- **green:** for genes with log2R changing homodirectionally in two condition.
- **yellow:** for genes with log2R changing antidirectionally in two condition.

##### 3.4.6 xtail/fc_<contrast>.pdf

This figure shows the result of the differential expression at the two expression levels, where each gene is a dot whose position is determined by its log2 fold change (log2FC) of transcriptional level (mRNA log2FC), represented on the x-axis, and the log2FC of translational level (RPF log2FC), represented on the y-axis. The points will be color-coded with the pvalue final obtained with xtail (more significant p values having darker color)

- **blue:** for genes whos mRNA log2FC larger than 1 (transcriptional level).
- **red:** for genes whos RPF log2FC larger than 1 (translational level).
- **green:** for genes changing homodirectionally at both level.
- **yellow:** for genes changing antidirectionally at two levels.

#### 3.5 Metagene Analysis

Meta gene profiling analyses the distribution of mapped reads around the start codon. Moreover for Ribo-seq it is expected that the ribosome protects a specific range of read lengths, often typical for the investigated group of organisms, from digestion by nuclease. These reads should show a typical peak around the start codon which corresponds to the high frequency that ribosomes are bound there. We output and plot the meta gene profiling for each individual fragment length as a quality control for the Ribo-seq protocol. If the distribution for all read lengths is untypical, arresting the ribosomes failed.

##### 3.5.1 <accession>_Z.Y_profiling.xlsx/tsv

The table shows for a range of specific read lengths, how many reads on average over all start codons in the genome have been mapped per nucleotide. The nucleotides range from 100 nucleotides upstream of the start codon to 399 nucleotides downstream. The read counts are either raw or normalized by average read count per nucleotide, for the range around the start codon. Moreover different single nucleotide mapping variants are considered, where only the 5', 3' or centered region of the read is counted.

##### 3.5.2 <accession>_Z.Y_profiling.pdf

#### 3.6 Additional output

##### 3.6.1 samples.xlsx

An excel representation of the input sample file.

##### 3.6.2 manual.pdf

A PDF format file giving some explanations about the output files, contained in the final result report.

##### 3.6.3 overview.xlsx

An overview table containing all information gathered from the prediction tools and differential expression analysis. The contents of this table change depending on which *options* are set. The overview table for the default workflow will contain annotation. reparation, deepribo and differential expression output.

| Column name | Description |
|---|---|
| Genome | The genome accession identifier. |
| Start | The start position of the ORF. |
| Stop | The stop position of the ORF. |
| Strand | The strand of the ORF. (+/-) |
| Locus_tag | The locus tag of ORF. (if not novel) |
| Name | The name of the ORF. (if not novel) |
| Gene_name | The name of the ORFs associated gene feature. (if not novel) |
| Length | The length of the ORF. |
| Codon_count | The number of codons in the ORF. (length / 3) |
| <method>-<condition>-<replicate>_TE | The translational efficiency for the given sample. |
| <method>-<condition>-<replicate>_rpkm | The RPKM for the given sample. (ReadsPerKilobaseMillion) |
| Start_codon | The start codon of the annotated feature. |
| Stop_codon | The stop codon of the annotated feature. |
| Nucleotide_seq | The nucleotide sequence of the annotated feature. |
| Aminoacid_seq | The amino acid sequence of the annotated feature. |
| Evidence | The tool and sample this ORF was predicted in (only for predicted ORFs) |
| Reparation_probability | The probability of this ORF. (only available for reparation predictions) |
| Deepribo_rank | The deepribo rank for this ORF. (only available for deepribo predictions) |
| Deepribo_score | The score the deepribo rank is based on. |
| riborex_pvalue | The pvalue (determined by riborex) |
| riborex_pvalue_adjusted | The adjusted pvalue (determined by riborex) |
| riborex_log2FC | The log2FC (determined by riborex) |
| xtail_pvalue | The pvalue (determined by xtail) |
| xtail_pvalue_adjusted | The adjusted pvalue (determined by xtail) |
| xtail_log2FC | The log2FC (determined by xtail) |

CHAPTER

FOUR

### EXAMPLE WORKFLOW

The retrieval of input files and running the workflow locally and on a server cluster via a queuing system is demonstrated using an example with data available from NCBI. This dataset is available under the accession number `PRJNA379630`.

**Note:** In this tutorial, we will show the basic functionalities of our workflow, for information about additional options please refer to: *workflow-configuration*.

**Note:** Ensure that you have `miniconda3` installed and a `snakemake environment` set-up. Please refer to the *overview* for details on the installation.

#### 4.1 Setup

First of all, we start by creating the project directory and changing to it.

```
mkdir project
cd project
```

We then download the latest version of HRIBO into the newly created project folder and unpack it.

```
wget https://github.com/RickGelhausen/HRIBO/archive/1.4.0.tar.gz
tar -xzf 1.4.0.tar.gz; mv HRIBO-1.4.0 HRIBO; rm 1.4.0.tar.gz;
```

#### 4.2 Retrieve and prepare input files

Before starting the workflow, we have to acquire and prepare several input files. These files are the annotation file, the genome file, the fastq files, the configuration file and the sample sheet.

##### 4.2.1 Annotation and genome files

First, we want to retrieve the annotation file and the genome file. In this case, we can find both on NCBI using the accession number `NC_002516.2`.

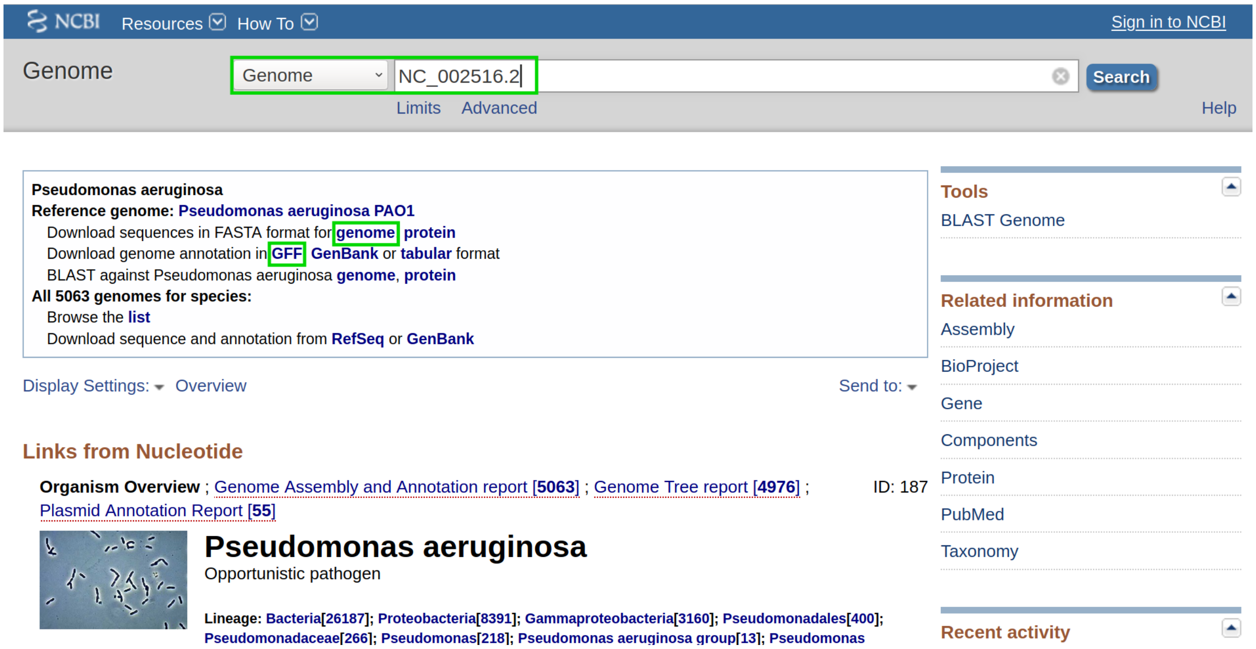

On this page, we can directly retrieve both files by clicking on the according download links next to the file descriptions. Alternatively, you can directly download them using the following commands:

```
wget ftp://ftp.ncbi.nlm.nih.gov/genomes/all/GCF/000/006/765/GCF_000006765.1_ASM676v1/
↪GCF_000006765.1_ASM676v1_genomic.gff.gz
wget ftp://ftp.ncbi.nlm.nih.gov/genomes/all/GCF/000/006/765/GCF_000006765.1_ASM676v1/
↪GCF_000006765.1_ASM676v1_genomic.fna.gz
```

Then, we unpack and rename both files.

```
gunzip GCF_000006765.1_ASM676v1_genomic.gff.gz && mv GCF_000006765.1_ASM676v1_genomic.
↪gff annotation.gff
gunzip GCF_000006765.1_ASM676v1_genomic.fna.gz && mv GCF_000006765.1_ASM676v1_genomic.
↪fna genome.fa
```

##### 4.2.2 .fastq files

Next, we want to acquire the fastq files. The fastq files are available under the accession number `PRJNA379630` on NCBI. The files have to be downloaded using the Sequence Read Archive (SRA). There are multiple ways of downloading files from SRA as explained here.

As we already have conda installed, the easiest way is to install the *sra-tools*:

```
conda create -n sra-tools -c bioconda -c conda-forge sra-tools pigz
```

This will create a conda environment containing the sra-tools. Using these, we can simply pass the SRA identifiers and download the data:

```
conda activate sra-tools;
fasterq-dump SRR5356908; pigz -p 2 SRR5356908.fastq; mv SRR5356908.fastq.gz RNA-PAO1-
↪gly-1.fastq.gz;
fasterq-dump SRR5356907; pigz -p 2 SRR5356907.fastq; mv SRR5356907.fastq.gz RIBO-PAO1-
↪gly-1.fastq.gz;
conda deactivate;
```

**Note:** Due to the runtime of several tools, especially the mapping by `segemehl`, this tutorial only uses one condition and replicate. If available, it is advisable to use as many replicates as possible.

**Warning:** If you have a bad internet connection, this step might take some time. If you prefer, you can also use your own `.fastq` files. But ensure that you use the correct annotation and genome files and that you compress them in `.gz` format (using gzip, pigz, etc . . . )

This will download compressed files for each of the required `.fastq` files. We will move them into a folder called `fastq`.

```
mkdir fastq;
mv *.fastq.gz fastq;
```

##### 4.2.3 Sample sheet and configuration file

Finally, we will prepare the configuration file (`config.yaml`) and the sample sheet (`samples.tsv`). We start by copying templates for both files from the `HRIBO/templates/` into the `HRIBO/` folder.

```
cp HRIBO/templates/samples.tsv HRIBO/
```

The sample file looks as follows:

| method | condition | replicate | fastqFile |
|---|---|---|---|
| RIBO | A | 1 | fastq/RIBO-A-1.fastq.gz |
| RIBO | A | 2 | fastq/RIBO-A-2.fastq.gz |
| RIBO | B | 1 | fastq/RIBO-B-1.fastq.gz |
| RIBO | B | 2 | fastq/RIBO-B-2.fastq.gz |
| RNA | A | 1 | fastq/RNA-A-1.fastq.gz |
| RNA | A | 2 | fastq/RNA-A-2.fastq.gz |
| RNA | B | 1 | fastq/RNA-B-1.fastq.gz |
| RNA | B | 2 | fastq/RNA-B-2.fastq.gz |

**Note:** When using your own data, use any editor (vi(m), gedit, nano, atom, . . . ) to customize the sample sheet.

**Warning: Please ensure not to replace any tabulator symbols with spaces while changing this file.**

We will rewrite this file to fit the previously downloaded *.fastq.gz* files.

| method | condition | replicate | fastqFile |
|---|---|---|---|
| RIBO | GLY | 1 | fastq/RIBO-PAO1-gly-1.fastq.gz |
| RNA | GLY | 1 | fastq/RNA-PAO1-gly-1.fastq.gz |

Next, we are going to set up the `config.yaml`.

```
cp HRIBO/templates/config.yaml HRIBO/
```

This file contains the following variables:

- **adapter:** Specify the adapter sequence to be used. In our case this would be *AGATCGGAAGAGCACACGTCT-GAACTCCAGTCAC*
- **samples:** The location of the sample sheet created in the previous step.
- **alternativestartcodons:** Specify a comma separated list of alternative start codons.
- **differentialexpression:** Specify whether you want to activate differential expresssion analysis. ("yes/no")
- **deepribo:** Specify whether you want to activate deepribo ORF prediction. ("yes/no")

In our example, this will lead to the following `config.yaml` file:

```
adapter: "AGATCGGAAGAGCACACGTCTGAACTCCAGTCAC"
samples: "HRIBO/samples.tsv"
alternativestartcodons: "GTG,TTG"
# Differential expression: on / off
differentialexpression: "off"
# Deepribo predictions: on / off
deepribo: "off"
```

#### 4.3 Running the workflow

Now that all the required files are prepared, we can start running the workflow, either locally or in a cluster environment.

> **Warning:** before you start using `snakemake` remember to activate the environment first.

```
conda activate snakemake
```

##### 4.3.1 Run the workflow locally

Use the following steps when you plan to execute the workflow on a single server, cloud instance or workstation.

> **Warning:** Please be aware that some steps of the workflow require a lot of memory or time, depending on the size of your input data. To get a better idea about the memory consumption, you can have a look at the provided `sge.yaml` or `torque.yaml` files.

Navigate to the project folder containing your annotation and genome files, as well as the HRIBO folder. Start the workflow locally from this folder by running:

```
snakemake --use-conda -s HRIBO/Snakefile --configfile HRIBO/config.yaml --directory $
↪{PWD} -j 10 --latency-wait 60
```

This will start the workflow locally.

- `--use-conda`: instruct snakemake to download tool dependencies from conda.
- `-s`: specifies the Snakefile to be used.
- `--configfile`: specifies the config file to be used.
- `--directory`: specifies your current path.
- `-j`: specifies the maximum number of cores snakemake is allowed to use.
- `--latency-wait`: specifies how long (in seconds) snakemake will wait for filesystem latencies until declaring a file to be missing.

##### 4.3.2 Run Snakemake in a cluster environment

Use the following steps if you are executing the workflow via a queuing system. Edit the configuration file `<cluster>.yaml` according to your queuing system setup and cluster hardware.

Navigate to the project folder on your cluster system. Start the workflow from this folder by running (The following system call shows the usage with Grid Engine.):

```
snakemake --use-conda -s HRIBO/Snakefile --configfile HRIBO/config.yaml --directory $
↪{PWD} -j 20 --cluster-config HRIBO/sge.yaml
```

**Note:** Ensure that you use an appropriate `<cluster>.yaml` for your cluster system. We provide one for `SGE` and `TORQUE` based systems.

##### 4.3.3 Example: Run Snakemake in a cluster environment

**Warning: Be advised that this is a specific example, the required options may change depending on your system.**

We ran the tutorial workflow in a cluster environment, specifically a TORQUE cluster environment. Therefore, we created a bash script `torque.sh` in our project folder.

```
vi torque.sh
```

**Note:** Please note that all arguments enclosed in <> have to be customized. This script will only work if your cluster uses the TORQUE queuing system.

We proceeded by writing the queuing script:

```
#!/bin/bash
#PBS -N <ProjectName>
#PBS -S /bin/bash
```

(continues on next page)

(continued from previous page)

```
#PBS -q "long"
#PBS -d <PATH/ProjectFolder>
#PBS -l nodes=1:ppn=1
#PBS -o <PATH/ProjectFolder>
#PBS -j oe
cd <PATH/ProjectFolder>
source activate HRIBO
snakemake --latency-wait 600 --use-conda -s HRIBO/Snakefile --configfile HRIBO/config.
↪yaml --directory ${PWD} -j 20 --cluster-config HRIBO/torque.yaml --cluster "qsub -N
↪{cluster.jobname} -S /bin/bash -q {cluster.qname} -d <PATH/ProjectFolder> -l
↪{cluster.resources} -o {cluster.logoutputdir} -j oe"
```

We then simply submitted this job to the cluster:

```
qsub torque.sh
```

Using any of the presented methods, this will run the workflow on the tutorial dataset and create the desired output files.

#### 4.4 Results

The last step will be to aggregate all the results once the workflow has finished running. In order to do this, we provided a script in the scripts folder of HRIBO called `makereport.sh`.

```
bash HRIBO/scripts/makereport.sh <reportname>
```

Running this will create a folder where all the results are collected from the workflows final output, it will additionally create compressed file in `.zip` format.

**Note:** A detailed explanation of the result files can be found in the *result section*.

**Note:** The final result of this example workflow, can be found here .

#### 4.5 References

CHAPTER

FIVE

### EXTENDED WORKFLOW

**Warning:** This tutorial shows a full run of the workflow with all options activated. For testing, we ran this example on a TORQUE cluster system and locally on a large cloud instance. The data is likely too large for running locally on an average laptop.

We show a run of the full workflow, including deepribo predictions and differential expression analysis, on data available from NCBI. For this purpose, we use a `salmonella enterica` dataset available under the accession number `PRJNA421559` [PGAR19].

**Warning:** Ensure that you have `miniconda3` and `singularity` installed and a `snakemake environment` set-up. Please refer to the *overview* for details on the installation.

#### 5.1 Setup

First of all, we start by creating the project directory and changing to it. (you can choose any directory name)

```
mkdir project
cd project
```

We then download the latest version of HRIBO into the newly created project folder and unpack it.

```
wget https://github.com/RickGelhausen/HRIBO/archive/1.4.0.tar.gz
tar -xzf 1.4.0.tar.gz; mv HRIBO-1.4.0 HRIBO; rm 1.4.0.tar.gz;
```

#### 5.2 Retrieve and prepare input files

Before starting the workflow, we have to acquire and prepare several input files. These files are the annotation file, the genome file, the fastq files, the configuration file and the sample sheet.

##### 5.2.1 Annotation and genome files

First, we want to retrieve the annotation file and the genome file. In this case, we can find both on NCBI using the accession number `NC_016856.1`.

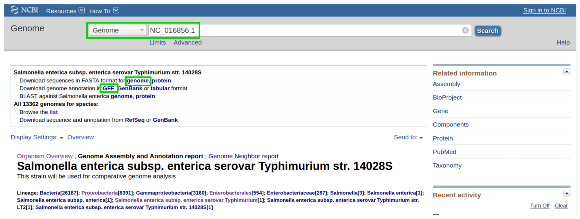

**Note:** Ensure that you download the annotation for the correct strain `str. 14028S`.

On this page, we can directly retrieve both files by clicking on the according download links next to the file descriptions. Alternatively, you can directly download them using the following commands:

```
wget ftp://ftp.ncbi.nlm.nih.gov/genomes/all/GCF/000/022/165/GCF_000022165.1_ASM2216v1/
↪GCF_000022165.1_ASM2216v1_genomic.gff.gz
wget ftp://ftp.ncbi.nlm.nih.gov/genomes/all/GCF/000/022/165/GCF_000022165.1_ASM2216v1/
↪GCF_000022165.1_ASM2216v1_genomic.fna.gz
```

Then, we unpack and rename both files.

```
gunzip GCF_000022165.1_ASM2216v1_genomic.gff.gz && mv GCF_000022165.1_ASM2216v1_
↪genomic.gff annotation.gff
gunzip GCF_000022165.1_ASM2216v1_genomic.fna.gz && mv GCF_000022165.1_ASM2216v1_
↪genomic.fna genome.fa
```

##### 5.2.2 .fastq files

Next, we want to acquire the fastq files. The fastq files are available under the accession number `PRJNA421559` on NCBI. The files have to be downloaded using the Sequence Read Archive (SRA). There are multiple ways of downloading files from SRA as explained here.

As we already have `conda` installed, the easiest way is to install the `sra-tools`:

```
conda create -n sra-tools -c bioconda -c conda-forge sra-tools pigz
```

This will create a conda environment containing the `sra-tools` and `pigz`. Using these, we can simply pass the SRA identifiers and download the data:

```
conda activate sra-tools;
fasterq-dump SRR6359966; pigz -p 2 SRR6359966.fastq; mv SRR6359966.fastq.gz RIBO-WT-1.
↪fastq.gz
fasterq-dump SRR6359967; pigz -p 2 SRR6359967.fastq; mv SRR6359967.fastq.gz RIBO-WT-2.
↪fastq.gz
fasterq-dump SRR6359974; pigz -p 2 SRR6359974.fastq; mv SRR6359974.fastq.gz RNA-WT-1.
↪fastq.gz
fasterq-dump SRR6359975; pigz -p 2 SRR6359975.fastq; mv SRR6359975.fastq.gz RNA-WT-2.
↪fastq.gz
fasterq-dump SRR6359970; pigz -p 2 SRR6359970.fastq; mv SRR6359970.fastq.gz RIBO-csrA-
↪1.fastq.gz
fasterq-dump SRR6359971; pigz -p 2 SRR6359971.fastq; mv SRR6359971.fastq.gz RIBO-csrA-
↪2.fastq.gz
```

(continues on next page)

(continued from previous page)

```
fasterq-dump SRR6359978; pigz -p 2 SRR6359978.fastq; mv SRR6359978.fastq.gz RNA-csrA-
↪1.fastq.gz
fasterq-dump SRR6359979; pigz -p 2 SRR6359979.fastq; mv SRR6359979.fastq.gz RNA-csrA-
↪2.fastq.gz
conda deactivate;
```

**Note:** we will use two conditions and two replicates for each condition. There are 4 replicates available for each condition, we run it with two as this is just an example. If you run an analysis always try to use as many replicates as possible.

**Warning:** If you have a bad internet connection, this step might take some time. It is advised to run this workflow on a cluster or cloud instance.

This will download compressed files for each of the required `.fastq` files. We will move them into a folder called `fastq`.

```
mkdir fastq;
mv *.fastq.gz fastq;
```

##### 5.2.3 Sample sheet and configuration file

Finally, we will prepare the configuration file (`config.yaml`) and the sample sheet (`samples.tsv`). We start by copying templates for both files from the `HRIBO/templates/` into the `HRIBO/` folder.

```
cp HRIBO/templates/samples.tsv HRIBO/
```

The sample file looks as follows:

| method | condition | replicate | fastqFile |
|---|---|---|---|
| RIBO | A | 1 | fastq/RIBO-A-1.fastq.gz |
| RIBO | A | 2 | fastq/RIBO-A-2.fastq.gz |
| RIBO | B | 1 | fastq/RIBO-B-1.fastq.gz |
| RIBO | B | 2 | fastq/RIBO-B-2.fastq.gz |
| RNA | A | 1 | fastq/RNA-A-1.fastq.gz |
| RNA | A | 2 | fastq/RNA-A-2.fastq.gz |
| RNA | B | 1 | fastq/RNA-B-1.fastq.gz |
| RNA | B | 2 | fastq/RNA-B-2.fastq.gz |

**Note:** When using your own data, use any editor (vi(m), gedit, nano, atom, . . . ) to customize the sample sheet.

**Warning: Please ensure not to replace any tabulator symbols with spaces while changing this file.**

We will rewrite this file to fit the previously downloaded *.fastq.gz* files.

| method | condition | replicate | fastqFile |
|---|---|---|---|
| RIBO | WT | 1 | fastq/RIBO-WT-1.fastq.gz |
| RIBO | WT | 2 | fastq/RIBO-WT-2.fastq.gz |
| RIBO | csrA | 1 | fastq/RIBO-csrA-1.fastq.gz |
| RIBO | csrA | 2 | fastq/RIBO-csrA-2.fastq.gz |
| RNA | WT | 1 | fastq/RNA-WT-1.fastq.gz |
| RNA | WT | 2 | fastq/RNA-WT-2.fastq.gz |
| RNA | csrA | 1 | fastq/RNA-csrA-1.fastq.gz |
| RNA | csrA | 2 | fastq/RNA-csrA-1.fastq.gz |

Next, we are going to set up the `config.yaml`.

```
cp HRIBO/templates/config.yaml HRIBO/
```

This file contains the following variables:

- **adapter:** Specify the adapter sequence to be used. In our case this would be *CTGTAGGCACCATCAAT*
- **samples:** The location of the sample sheet created in the previous step.
- **alternativestartcodons:** Specify a comma separated list of alternative start codons.
- **differentialexpression:** Specify whether you want to activate differential expresssion analysis. ("yes/no")
- **deepribo:** Specify whether you want to activate deepribo ORF prediction. ("yes/no")

In our example, this will lead to the following `config.yaml` file:

```
adapter: "CTGTAGGCACCATCAAT"
samples: "HRIBO/samples.tsv"
alternativestartcodons: "GTG,TTG"
# Differential expression: on / off
differentialexpression: "on"
# Deepribo predictions: on / off
deepribo: "on"
```

#### 5.3 Running the workflow

Now that all the required files are prepared, we can start running the workflow, either locally or in a cluster environment.

**Warning:** if you have problems running `deepribo`, please refer to *Activating DeepRibo*.

**Warning:** before you start using `snakemake` remember to activate the environment first.

```
conda activate snakemake
```

##### 5.3.1 Run the workflow locally

Use the following steps when you plan to execute the workflow on a single server, cloud instance or workstation.

> **Warning:** Please be aware that some steps of the workflow require a lot of memory or time, depending on the size of your input data. To get a better idea about the memory consumption, you can have a look at the provided `sge.yaml` or `torque.yaml` files.

Navigate to the project folder containing your annotation and genome files, as well as the HRIBO folder. Start the workflow locally from this folder by running:

```
snakemake --use-conda --use-singularity --singularity-args " -c " --greediness 0 -s␣
↪HRIBO/Snakefile --configfile HRIBO/config.yaml --directory ${PWD} -j 10 --latency-
↪wait 60
```

This will start the workflow locally.

- `--use-conda`: instruct snakemake to download tool dependencies from conda.
- `-s`: specifies the Snakefile to be used.
- `--configfile`: specifies the config file to be used.
- `--directory`: specifies your current path.
- `-j`: specifies the maximum number of cores snakemake is allowed to use.
- `--latency-wait`: specifies how long (in seconds) snakemake will wait for filesystem latencies until declaring a file to be missing.

##### 5.3.2 Run Snakemake in a cluster environment

Use the following steps if you are executing the workflow via a queuing system. Edit the configuration file `<cluster>.yaml` according to your queuing system setup and cluster hardware.

Navigate to the project folder on your cluster system. Start the workflow from this folder by running (The following system call shows the usage with Grid Engine):

```
snakemake --use-conda --use-singularity --singularity-args " -c " -s HRIBO/Snakefile -
↪-configfile HRIBO/config.yaml --directory ${PWD} -j 20 --cluster-config HRIBO/sge.
↪yaml
```

> **Note:** Ensure that you use an appropriate `<cluster>.yaml` for your cluster system. We provide one for `SGE` and `TORQUE` based systems.

**Note:** The final result of this example workflow, can be found here .

#### 5.5 References

### CHAPTER SIX

### FREQUENTLY ASKED QUESTIONS

#### 6.1 Q: When using singularity I get `ERROR : Failed to set effective UID to 0.`

Prior to the installation of singularity change `with_suid=1` to `with_suid=0` in the mconfig file in the singularity folder. This should not be necessary for newer versions of singularity.
